# STARR-CRAAVT: A platform to design cell type-specific regulatory elements for next-generation gene therapy

**DOI:** 10.1101/2025.06.04.657008

**Authors:** Robert Becker, Priyanka Choudhury, Martin Oti, Youssef Fouani, Benjamin Strobel, Stephanie Ketterer, Simon Rumpel, John Park, Sebastian Kreuz, Udo Maier, Stefan Michelfelder, Christian Schön

## Abstract

Precise control of transgene expression through novel enhancer-promoter combinations is a promising strategy for advancing gene therapy mediated by adeno-associated virus (AAV). We present STARR-CRAAVT, a novel STARR-Seq-derived platform to screen for enhancers in the AAV context. Using HepG2 and HaCaT cells as screening models, we applied STARR-CRAAVT for the identification of cell type-specific enhancers. We integrated epigenetic datasets into an *in-silico* library of putative HepG2 enhancers and captured corresponding fragments from genomic DNA. The fragments were processed into AAV libraries and applied to HepG2 and HaCaT cells. STARR-CRAAVT analysis revealed a selective activity of the libraries confirming the HepG2-directed *in-silico* design and the specificity of single enhancer-promoter combinations could be validated using luciferase reporter assays. In addition, we scrutinized the impact of key experimental parameters on enhancer identification and found that the used promoter type had significant influence on the ability of candidates to act as enhancers. Furthermore, switching the location of identified enhancers in reporter assays revealed that the level of enhancer activity is highly dependent on the position in the AAV genome. Taken together, our study yields novel insights into enhancer function and demonstrates that STARR-CRAAVT can be employed to identify cell type-specific enhancers, highlighting promoter preference and enhancer positioning as key considerations for enhancer screening campaigns.

## INTRODUCTION

Recombinant adeno-associated virus (rAAV) is the most widely used vector for gene therapy *in vivo* with currently eight approved rAAV-based gene therapy products and over 300 ongoing clinical trials in the US (clinicaltrials.gov, assessed on March 31^st^ 2025). However, to fully harness the potential of rAAVs and unlock new target cell types and diseases, further enhancements of specificity and potency are needed. Efforts to enhance rAAVs focus mainly on the engineering of 1) novel rAAV capsids to efficiently and specifically deliver the genetic material and 2) the rAAV genome to regulate transgene expression in the desired spatiotemporal manner. While capsid engineering has been extensively explored using rational design approaches as well as high-throughput screenings (Becker et al. 2022; Pupo et al. 2022), the resources and attention for rAAV genome engineering are only beginning to catch up.

In its most basic configuration, the single-stranded rAAV genome consists of the promoter-driven therapeutic transgene and two inverted terminal repeats (ITRs) flanking the transgene cassette, which are required for AAV packaging. Promising rAAV genome engineering approaches include introduction of miRNA binding sites into the transgene 3’ UTR, riboswitches, codon optimization of the transgene coding sequence, and the mutation of the ITRs to create self-annealing AAV genomes, which results in a faster expression kinetic by omitting second strand synthesis (McCarty 2008; Domenger and Grimm 2019; Strobel et al. 2020). However, one of the most potent strategies for expression control is the design of novel promoter regions. For this, promoters are combined with enhancer sequences, which confer additional strength and are often critical determinants of cell type-specific expression (Heinz et al. 2015; Jüttner et al. 2019). For clarity, in the following we will refer to these combinations of promoters and enhancers as *cis*-regulatory regions. Rational design of *cis*-regulatory regions derived from promoter and enhancer sequences of cell type-specific genes has been utilized successfully in gene therapy products, namely the various liver-specific derivates (HLP, LP1, ApoE/hAAT) in Roctavian (Bunting et al. 2018; McIntosh et al. 2013), Hemgenix (Nathwani et al. 2006) and Beqvez (George et al. 2017) for haemophilia therapy or the striated muscle-specific MHCK7 element in Elevidys for therapy of Duchenne’s muscular dystrophy (Mendell et al. 2020; Salva et al. 2007). Additionally, several rationally designed *cis*-regulatory regions have been developed for pre-clinical and clinical studies in other target organs, such as the eye (reviewed in (Buck and Wijnholds 2020)), the heart (reviewed in (Schröder et al. 2023)) and the CNS (Islam and Tom 2022; Nieuwenhuis et al. 2021). Yet, the identification of specific and potent promoter-enhancer combinations by rational design is eventually limited by the throughput of testing candidates in single AAVs and by the knowledge about enhancer-promoter compatibility, which is an ongoing research focus in the field of transcriptional regulation (Yang and Hansen 2024; Martinez-Ara et al. 2022; Bergman et al. 2022).

A powerful alternative to rational design approaches are massive parallel reporter assays (MPRAs), functional high-throughput screens that infer promoter/enhancer activity of up to millions of candidates by quantifying reporter transcripts via next-generation sequencing (NGS). One of these MPRAs is STARR-Seq (self-transcribing active regulatory region sequencing), a barcode-independent screening for enhancers (Arnold et al. 2013). In STARR-Seq, candidate libraries are placed downstream of a promoter resulting in transcription of the candidate sequences. Enhancers are expected to activate the promoter, thereby increasing their own transcription. STARR-Seq has been applied in the AAV context in two previous studies, distinguished by their candidate library design. Initial work demonstrating the feasibility of combining STARR-Seq with AAV used PCR to generate a library with hundreds of candidates from the mouse genome (Lambert et al. 2021). In a subsequent study aiming to identify enhancers in the adult mouse brain, we used BACs covering large genomic regions (∼ 3Mb total) containing brain-specific genes (Chan et al. 2023). With this strategy, we were able to achieve delivery of ∼500,000 distinct candidates, an unprecedented library complexity for an AAV-based MPRA. Notably, both studies identified novel, potent regulatory elements driving reporter transgene expression *in vivo*, demonstrating that AAV-based STARR-Seq is a powerful tool for enhancer/promoter engineering.

Here, we combine for the first time rAAV-based STARR-Seq with a capture-based library creation similar to the CapSTARR-Seq approach (Vanhille et al. 2015), which allows precise and flexible incorporation of genomic fragments covering *in-silico*-nominated target sequences into the screening library while maintaining high library complexity. We term this screening platform STARR-CRAAVT (**<u>s</u>**elf-**t**ranscribing **<u>a</u>**ctive **<u>r</u>**egulatory **<u>r</u>**egion sequencing of **<u>c</u>**aptured libraries for **<u>rAAV</u>** gene **t**herapy). We performed an *in-vitro* screening campaign, addressing relevant applications and questions regarding STARR-CRAAVT with the aim to facilitate the use of this technology in the design of next-generation gene therapy vectors. Importantly, we show that comparative analysis of STARR-CRAAVT results can be used to identify cell type-specific enhancer elements from a given library. Furthermore, we find that the use of identified enhancers in a building block principle (e.g. in different vector designs) is limited by promoter preferences of enhancer candidates as well as the effect of relative positioning on enhancer activity. Taken together, our findings represent novel insights into enhancer function and will help to guide the design of future screening campaigns for regulatory elements in the AAV context.

## RESULTS

### Generation of the screening libraries

A main objective of this study was to determine whether STARR-CRAAVT can be employed in a comparative manner to identify cell type-specific enhancers that could be used in the design of cell-specific therapeutic AAVs. For this proof-of-concept study, we chose HepG2 cells, a hepatocellular carcinoma (HCC) cell line that has been used extensively as an MPRA model cell line (Klein et al. 2020, 2018; Lee et al. 2020), as the target cell type of our screen. In addition, we used HaCaT cells of keratinocyte origin as the “off-target” cell type, to enable comparative screening. To combine AAV-based STARR-Seq with a capture-based screening library and enable STARR-CRAAVT, we first designed an *in-silico* library of candidate genomic regions. We decided to tailor this library according to our primary target cell type HepG2. To build an HepG2/HCC-focused candidate library, we re-analysed, integrated, and filtered publicly available datasets (see Figure 1A for a graphic representation of the design strategy). Our starting point was a set of open chromatin regions derived from ATAC-Seq data of 17 HCCs from the TCGA database (Corces et al. 2018). We pruned this extensive list of 121,608 candidate regions by removing those overlapping transcription start sites to focus on potential enhancer elements. In a second step, we retained only regions marked by histone 3 lysine 27 acetylation (H3K27ac), a hallmark of active enhancers (Shlyueva et al. 2014), in HCC samples (Hlady et al. 2019). As HCC arises from hepatocytes, we decided to remove regions with high H3K27ac in normal hepatocytes (Moore et al. 2020), reasoning that this would focus the library more on the cancerous state of HCC/HepG2 cells. The resulting list contained 11,349 candidate regions (Figure 1A). As there was a substantial part of H3K27ac sites in HCC samples that did not overlap with ATAC-Seq peaks, we decided to supplement our candidate list with those H3K27 regions. After additional filtering steps to remove miRNAs and putative insulators, we arrived at the final list constituting 15,379 candidate regions (Figure 1A). Fragments covering the *in-silico* candidate regions were captured from sheared genomic DNA using the Agilent SureSelect^XT^ Target Enrichment System (Figure 1B). Briefly, candidate fragments were captured using biotinylated RNA oligos as bait and purified by streptavidin-coated magnetic beads (Gnirke et al. 2009). Purified fragments were subsequently bulk cloned into AAV transfer plasmids (Figure 1B).

**Figure 1.**
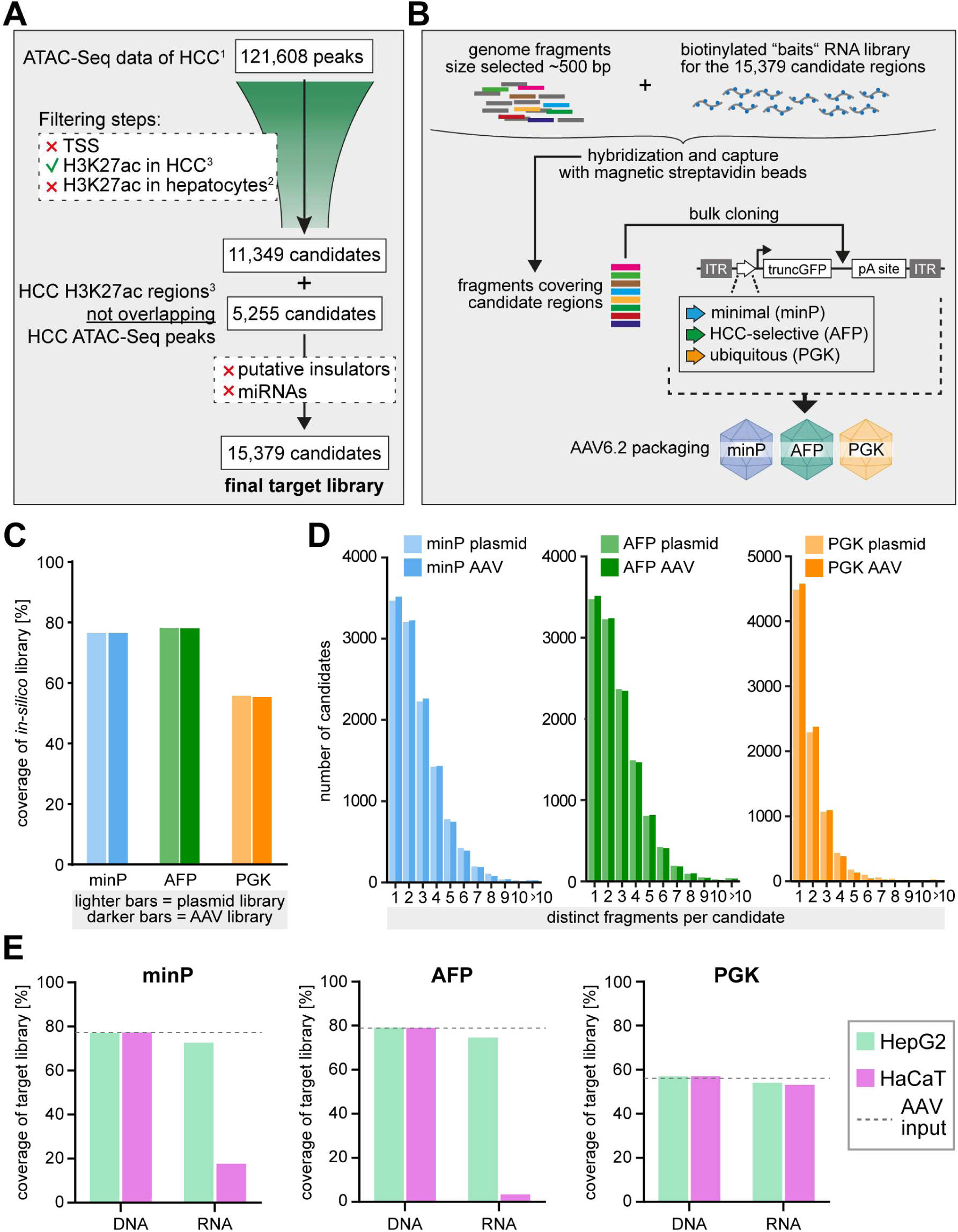
STARR-CRAAVT candidate library design and preparation of screening libraries. **(A)** Schematic overview of the design strategy used for the *in-silico* candidate library. **(B)** Generation of the screening libraries via the capture of genomic fragments, cloning, and AAV packaging. **(C)** Coverage of the *in-silico*-designed candidate library within the plasmid and virus libraries. **(D)** Distribution of distinct fragment counts per candidate in the plasmid and virus libraries. **(E)** Coverage of the *in-silico*-designed candidate library by fragments detected on DNA and RNA level in HepG2 or HaCaT cells after transduction with the respective AAV libraries. Dashed lines indicate coverage within the input AAV libraries from (C). HCC: hepatocellular carcinoma. ITR: inverted terminal repeat. truncGFP: truncated GFP. TSS: transcriptional start site.

Besides the identification of cell type-specific enhancers, a second objective of this study was to evaluate the promoter preference of candidates in our STARR-CRAAVT approach. To enable this, the captured candidate fragments were processed into three different AAV transfer plasmid libraries, each harbouring a different type of core promoter (Figure 1B): 1) a 32-bp minimal TATA-box promoter element (minP, previously referred to as “4.26” (Chan et al. 2023)), 2) a 276-bp promoter sequence of the fetal liver-specific alpha fetoprotein (AFP) gene, which has been shown to drive HCC-selective expression (Kanai et al. 1997) and 3) the 511-bp phosphoglycerate kinase 1 promoter (PGK) that is ubiquitously active and drives consistent expression across different mammalian cell types (Qin et al. 2010). The transfer plasmid libraries were packaged into AAV6.2 capsids, which have been shown to promote efficient transduction of HepG2 (Weinmann et al. 2022). Using our 50-AAV capsid panel, we confirmed that AAV6.2 transduces HaCaT cells as well (Figure S1). To assess the efficiency of the capture-cloning process and the impact of AAV packaging, we analysed the composition of the plasmid and AAV libraries by NGS. On plasmid level, our analysis revealed that minP and AFP candidate libraries contained approx. 32,000 distinct fragments that covered ∼77 and ∼79% of *in-silico* target regions, respectively. The PGK candidate library contained approx. 17,000 distinct fragments covering ∼57% of *in-silico* target regions (Figure 1C). The vast majority of candidates in all three libraries was covered by less than five distinct fragments (Figure 1D). Importantly, AAV packaging did not significantly impact the target region coverage nor the distribution of fragments per candidate, regardless of the promoter type (Figures 1C, 1D). Furthermore, analysis of normalized counts indicated a high consistency of target region abundance between plasmid and AAV libraries (Figure S2A; Pearson r: 0.89 (minP), 0.94 (AFP), and 0.98 (PGK)).

### STARR-CRAAVT screen

To perform STARR-CRAAVT, we transduced HepG2 and HaCaT cells with the three AAV enhancer libraries (minP, AFP, PGK) and isolated DNA and RNA after 72h. The transcriptional activity of a candidate in the screen (subsequently referred to as STARR-Seq activity) was quantified as the log2 fold change (log2FC) of RNA reads (i.e. expression level of the candidate) over DNA reads (i.e. input amount present in the cells). Before analysing STARR-Seq activities, we first assessed the transduction efficiency of each library by quantifying target region coverage on DNA level compared to the AAV input libraries. For all three libraries, we observed a target region coverage at input library levels in both, HepG2 and HaCaT cells, indicating delivery of all regions from the corresponding input libraries (Figure 1E). To assess overall library expression, we analysed the coverage of the target regions on RNA level in both cell types. In HepG2 cells, we observed a coverage close to input library levels for all three libraries, indicating a robust overall library expression, regardless of the promoter used. In contrast, coverage in HaCaT cells transduced with minP and AFP libraries were markedly lower compared to the corresponding input levels (Figure 1E). Only PGK library-transduced cells showed a coverage near input levels. These results indicate that the HepG2-selective activity of the AFP promoter is not affected globally by the candidate library. Furthermore, the lower minP-driven expression levels in HaCaT cells suggest that the overall library activity is weaker in those cells, confirming that our *in-silico* design strategy resulted in the generation of an HepG2/HCC-focused library. However, in combination with a potent ubiquitous promoter (i.e. PGK), the library can be expressed robustly in HaCaT cells as well. In summary, the coverage analyses indicate sufficient delivery and activity of the libraries to enable comparative STARR-CRAAVT analysis.

Classical STARR-Seq and CapSTARR-Seq approaches use plasmid-based delivery (Arnold et al. 2013; Vanhille et al. 2015). Therefore, prior to comparative STARR-CRAAVT analysis, we leveraged the advantage of an *in-vitro* system being amenable to different delivery methods and introduced the plasmid minP candidate library to HepG2 cells by transfection to assess whether the mode of delivery (plasmid vs. AAV) influences the STARR-Seq outcome (Figure S3). When comparing the STARR-Seq activities, we observed a good correlation between plasmid and AAV-based delivery (Pearson r = 0.763, p-value < 0.001), suggesting that the mode of delivery does not have a pronounced effect on overall candidate activity. Next, we aimed to estimate how the delivery mode would influence nomination of promising candidates, a main intention of high-throughput screens for regulatory element development. We compared the ranking of top performing candidates according to STARR-Seq activity between plasmid and AAV-based delivery (Figure S3). For ranking the 1,000 top performing candidates, we observed a moderate positive correlation (Spearman r = 0.428, p-value < 0.001). In comparison, for the 100 top performing candidates, a correlation between the rankings was notably lower (Spearman r = 0.197, p-value = 0.0226). Among the 100 top candidates, 57 were common to plasmid and AAV-delivered samples, albeit at different ranks. Four candidates were the top performing candidates in both conditions. Taken together, the mode of delivery did not have a pronounced impact on overall library activity. However, the ranking of candidates according to their STARR-Seq activity can change significantly based on the used modality, which should be considered when nominating top enhancer candidates for regulatory element development.

### STARR-CRAAVT can be used to identify cell type-specific enhancers

To test the utility of STARR-CRAAVT for identifying cell type-specific enhancers, we first compared the STARR-Seq activity of the enhancer libraries between HepG2 and HaCaT cells. The correlation of the activities in HepG2 and HaCaT cells was comparably low for minP and AFP AAV libraries (Pearson r = 0.537, p-value < 0.001 and r = 0.395, p-value < 0.001, respectively), while for the PGK AAV library we observed a higher correlation (Pearson r = 0.844, p-value < 0.001) (Figure 2A). This demonstrates that the activity profiles of the libraries change dependent on the cell type and suggests that STARR-CRAAVT can be used to detect cell type-specific differences in candidate activity. Next, we aimed to identify cell type-specific enhancers via comparative analysis of the STARR-CRAAVT outcome. To this end, we first determined the number of identified enhancers (defined here by a log2FC RNA/DNA > 1 and an adjusted p-value < 0.05) per library in each cell type. Comparing minP AAV library-transduced samples, we found substantially more enhancers in HepG2 (1401) than in HaCaT cells (331). The difference in identified enhancers was even more pronounced for the AFP AAV library. Specifically, 1982 enhancers were identified in HepG2 but only one in HaCaT cells. While a few more candidates exhibited a log2FC RNA/DNA > 1 in HaCaT cells, none of these reached significance. These results are again in line with the HepG2/HCC-focused library design strategy and suggest that only a small number of candidates can change the specificity of the AFP promoter. In contrast, the number of identified enhancers was similar between HepG2 (891) and HaCaT (952) for the PGK AAV library. Next, we quantified the fractions of cell type-specific enhancers for our target cell type HepG2, focusing on the data obtained with the minP and PGK AAV libraries (Fig. 2B). For the minP AAV library, we found ∼89% (1243/1401) of HepG2 enhancers to be specific for this cell type. In contrast, for the PGK AAV library, percentages of HepG2-specific enhancers were lower with ∼33% (291/891) of all HepG2 enhancers. To confirm the comparative AAV-STARR-Seq approach for the identification of cell type-specific enhancers, we tested a set HepG2-exclusive enhancers identified with minP in a single AAV setup using a luciferase reporter assay in HepG2 and HaCaT cells. Indeed, we could confirm significant enhancer activity for all enhancer sequences compared to a control stuffer sequence in HepG2 cells (Figure 2C). Furthermore, all tested sequences showed repressive behaviour compared to control when HaCaT cells were transduced (Figure 2C). Taken together, these results indicate that STARR-CRAAVT can be used in a comparative manner to identify cell type-specific enhancers. Yet, the ability of enhancers to drive expression in a cell type-specific fashion appears to vary significantly, dependent on the used promoter.

**Figure 2.**
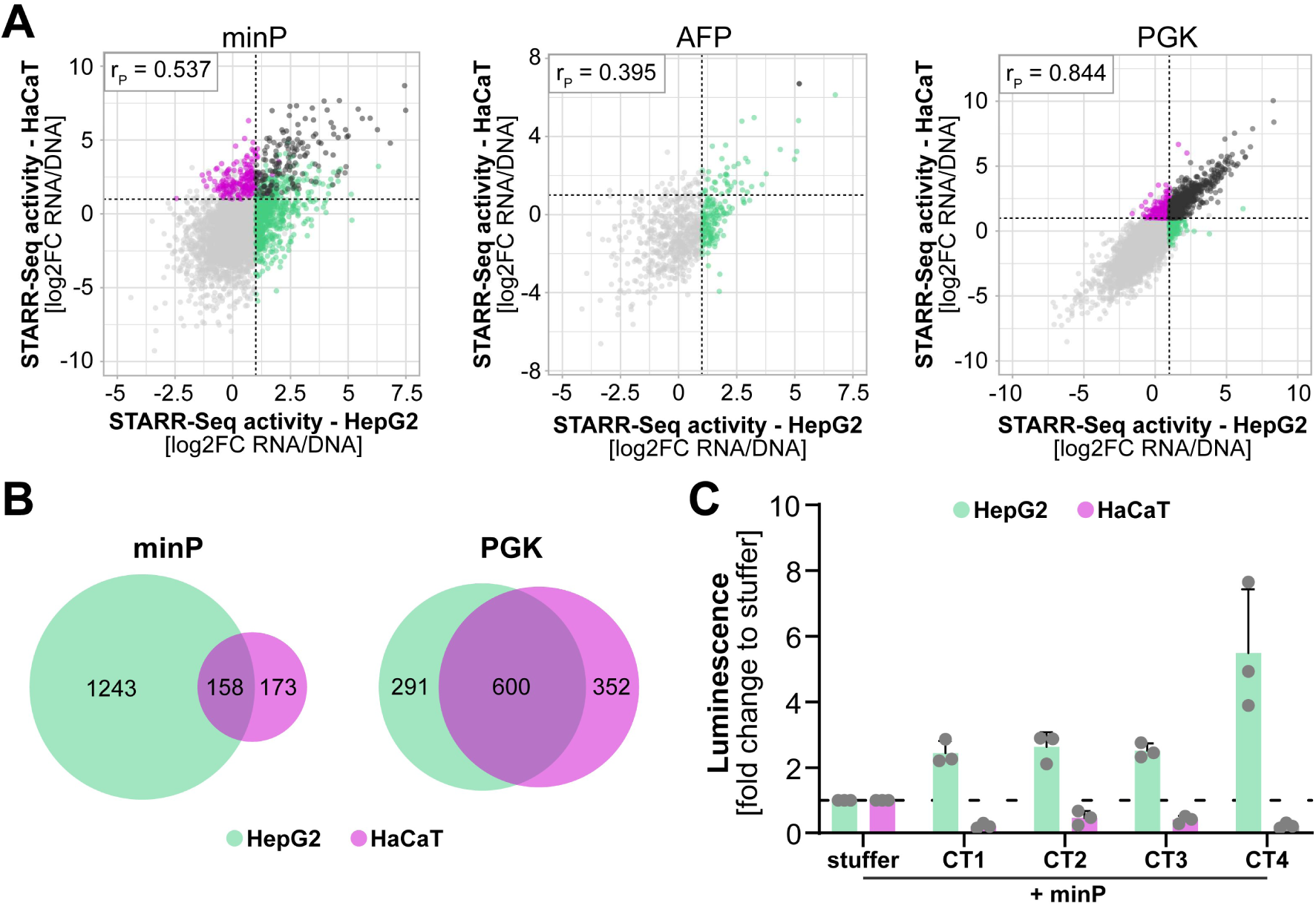
Comparative AAV-STARR-Seq identifies cell type-specific enhancers. **(A)** Scatterplots comparing STARR-Seq activity (measured as log2FC RNA/DNA) of candidate libraries between HepG2 and HaCaT cells. Graphs show only candidates for which a log2FC RNA/DNA could be calculated for both cell types. Dotted lines indicate the STARR-Seq activity cut-off for enhancers (log2FC RNA/DNA > 1). Colours indicate significant (p < 0.05) enhancer activity in HepG2 (●), HaCaT (●), or both (●). rP: Pearson correlation coefficient. **(B)** Venn diagrams showing the number of enhancers (log2FC RNA/DNA > 1, p < 0.05) per cell type in the respective candidate library. **(C)** Luciferase assay of selected HepG2-specific enhancer candidates (cell type-specific (CT) 1-4) tested as single AAV candidates together with minP. Activity is shown as fold change compared to activity of a neutral stuffer sequence (dashed line).

### Enhancer activity in AAV-STARR-Seq is highly promoter-dependent

A frequent strategy to rationally design regulatory elements for AAV gene therapy is the combination of promoters and enhancers in a building block principle (Domenger and Grimm 2019; Nathwani et al. 2006; Salva et al. 2007). Broader systematic studies of enhancer-promoter compatibility have yielded somewhat contradictory results to what extent this building block approach can be generalized (Bergman et al. 2022; Martinez-Ara et al. 2022; Yang and Hansen 2024). Whether identified enhancers can be employed in a modular fashion together with various promoters is also an important question for the use of STARR-CRAAVT for developing regulatory elements, as this would increase versatility and efficiency of enhancer screening campaigns. Therefore, we addressed this by comparing the STARR-CRAAVT outcomes in HepG2 cells, in which a good overall activity could be detected for all three libraries. Remarkably, correlation of STARR-Seq activities calculated as Pearson r was low between cells transduced with the different libraries (Figure 3A): 0.15 for minP library vs. AFP library, 0.08 for minP library vs. PGK library, and 0.1 for AFP library vs. PGK library (all p-value < 0.001). This suggests that overall activity of the candidate libraries in HepG2 is highly dependent on the used promoter. To further explore this, we compared the identified enhancers for each promoter-candidate library combination in HepG2 cells. As a pre-requisite, we assessed which candidates were present in all input AAV libraries, as only those would be eligible for a meaningful analysis of promoter preference. We found 6,482 out of 15,379 regions were covered by all three libraries, forming a robust basis for analysis of promoter preference (Figure S4). We then analysed which of those behaved as enhancers (log2FC > 1, adjusted p-value < 0.05) in transduced HepG2 cells, identifying 774 enhancers for the minP library, 1,070 enhancers for the AFP library, and 662 enhancers for PGK library (Figure 3B). Importantly, most enhancers were promoter-exclusive (∼66% for minP, ∼74% for AFP, ∼68% for PGK). In contrast, only 29 enhancers (∼0.4% of the common input library fragments) were found to act as enhancers with all three promoters (Figure 3B). Taken together, these findings demonstrate that in our AAV-STARR-Seq library setup enhancer activity of sequences is largely promoter-dependent.

**Figure 3.**
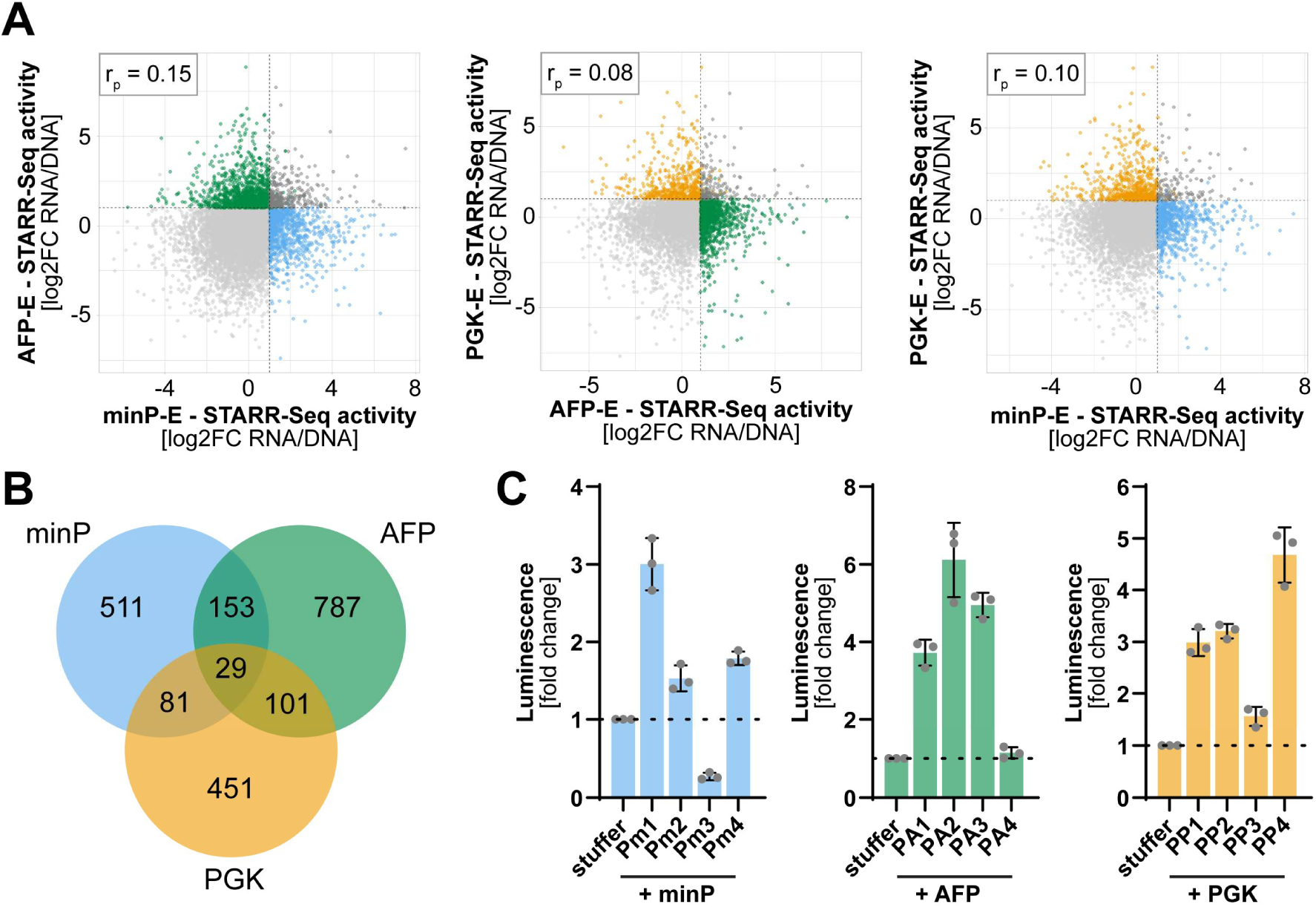
Enhancer activity is predominantly promoter-selective. **(A)** Scatterplots comparing candidate activities in HepG2 cells transduced with minP-E, AFP-E, and PGK-E as indicated in the axis labels. Activity was measured as log2FC RNA/DNA. Graphs show only candidates for which a log2FC RNA/DNA could be calculated for both cell types. Dotted lines indicate the STARR-Seq activity cut-off for enhancers (log2FC RNA/DNA > 1). Colours indicate significant (p < 0.05) enhancer activity with minP (●), AFP (●), PGK (●), or both promoters in each plot (●). rP: Pearson correlation coefficient. **(B)** Venn diagram showing the number of enhancers (log2FC RNA/DNA > 1, p < 0.05) per promoter in HepG2 cells. **(C)** Luciferase assays of selected enhancer/promoter combinations tested as single AAV constructs in HepG2 cells. Enhancers were chosen to be exclusive candidates from minP (Pm 1-4), AFP (PA 1-4) and PGK (PP 1-4) libraries. Activity is shown as fold change to a control stuffer sequence.

**Figure 4.**
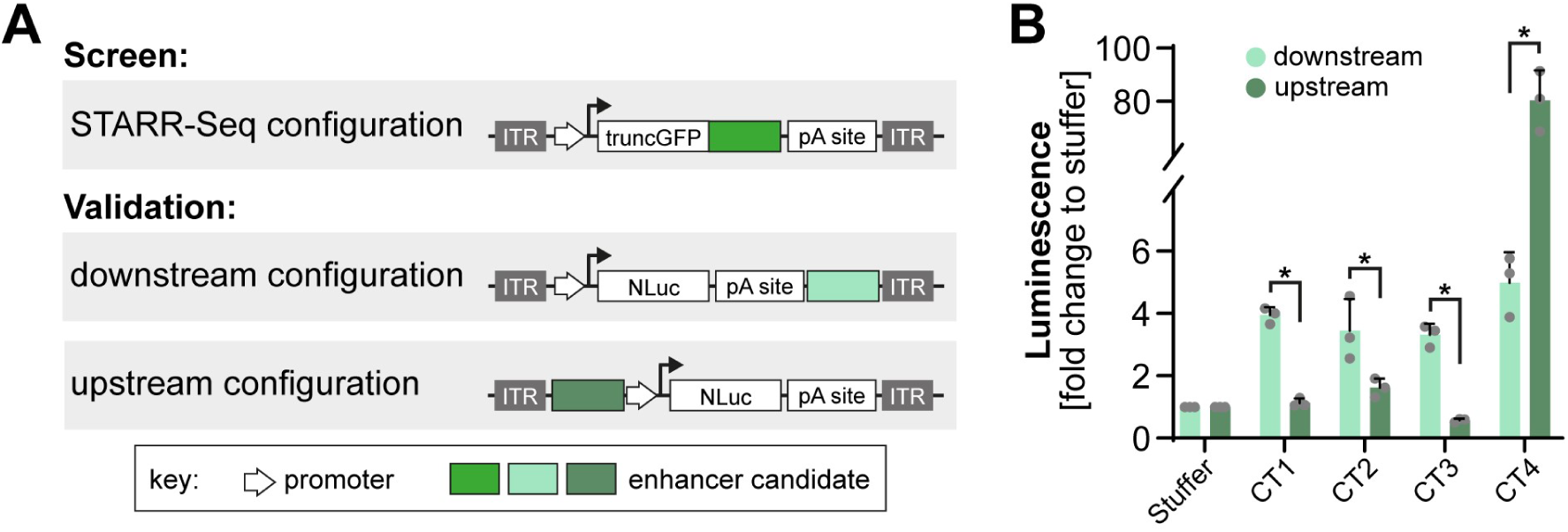
Enhancer activity is position-sensitive. **(A)** Schematic representation of the AAV expression constructs used in this study for screening and validation and their differences in enhancer candidate positioning. **(B)** Luciferase assay in HepG2 cells of the validated cell type-specific enhancers (CT1-CT4) combined with minP in the downstream and the upstream configuration. Activity is shown as fold change compared to a control stuffer sequence. *: p < 0.05.

To again confirm AAV-STARR-Seq results, we tested sets of promoter-specific enhancers in single AAV luciferase assays in HepG2 cells (Figure 3C). For minP and AFP, we could validate enhancer activity for 3 out of 4 tested sequences compared to a stuffer sequence, while for PGK all tested candidates showed significant enhancer activity. This further verifies the robustness of our AAV-STARR-Seq approach.

### Enhancer activity is position-sensitive

It is widely accepted that on a genomic level enhancers function is largely independent of their relative position to the promoter inside a certain distance (Panigrahi and O’Malley 2021). In most AAV expression cassettes, enhancers are placed directly upstream of the promoter controlling expression of an open reading frame (ORF). In contrast, in AAV-STARR-Seq and our corresponding reporter validation assays (Fig. 2C, 3C), enhancers are placed downstream of the ORF. To assess whether the position in the AAV expression cassette impacts enhancer activity, we tested the set of HepG2-specific enhancers that we had validated in the downstream configuration (Fig. 2C) by placing them upstream of the minP element and performing luciferase assays. Surprisingly, we found that enhancer activity varied substantially between the two configurations. For three out of four tested candidates, enhancer activity was either lost or significantly reduced when placed in the upstream configuration. In contrast, for the fourth candidate, activity was increased ∼16-fold compared to the downstream configuration. This indicates that in the context of the AAV genome and in contrast to the general assumption about enhancers (Thomas and Buecker 2023), positioning relative to the promoter can have a significant influence on enhancer activity/identity.

## DISCUSSION

Two recent studies in the mouse CNS demonstrated the potential of AAV-mediated STARR-Seq screening for identifying potent regulatory elements (Chan et al. 2023; Lambert et al. 2021). Here, we leveraged the strength of an *in-vitro* system as a highly controlled and scalable screening model to establish STARR-CRAAVT, a novel CapSTARR-Seq-inspired enhancer screening platform in the AAV context and pinpoint key parameters for the technology. We show that an *in-silico-* guided approach can be used to generate a cell type-focused candidate library and that AAV packaging does not significantly impact candidate library composition. Furthermore, we achieved delivery of the screening library to both, the target cell type HepG2 and the “off-target” cell type HaCaT, with a coverage at input levels enabling robust activity analysis. Using a capture approach to generate the screening library offers the advantage that a variety of regions from different genomic locations can be incorporated into the library based on defined criteria, for example open and active chromatin marks in a desired cell type. While this could in principle be achieved with synthesis-based library generation, our approach exhibits superior cost efficiency and scalability, exemplified here by library sizes of up to 32,000 fragments with a mean size of ∼400 bp. Therefore, STARR-CRAAVT is a promising platform for *cis*-regulatory region design for rAAV-mediated gene therapy Selective activation of transcription in specific cell types or cell states is among the most important criteria for an enhancer sequence to be used in next-generation rAAV vectors. As a proof-of-principle that STARR-CRAAVT is an adequate tool to identify cell type-specific enhancers, we applied our candidate library to two different cell types and performed comparative analysis of STARR-CRAAVT results. We identified enhancer candidates specifically driving expression in HepG2 but not HaCaT cells and confirmed the cell type specificity in single candidate luciferase assays. This insight will be valuable for future enhancer screening campaigns *in vivo* or in complex *in-vitro* models, which could combine STARR-CRAAVT with cell sorting or single cell-sequencing approaches, enabling comparative analysis on a multicellular level.

Promoter preferences of enhancer sequences remain a major area of research in the field of transcriptional control (Yang and Hansen 2024). Some studies report a high degree of enhancer-promoter specificity (Martinez-Ara et al. 2022), while others find enhancers to be largely promoter promiscuous (Bergman et al. 2022). These findings have been in part explained by different ways of analysis and model systems (i.e. mouse embryonic stem cells vs. human cell lines). Furthermore, at least in fruit flies enhancers regulating housekeeping genes differ from those regulating developmental genes in terms of promoter compatibility (Zabidi et al. 2015). In the context of AAV gene therapy vectors, knowledge about enhancer-promoter promiscuity is particularly useful to estimate whether an identified strong or specific enhancer can be used in a vector design-agnostic manner to tune expression. We find that in our STARR-CRAAVT approach, enhancer activity depends significantly on the promoter. Notably, we have tested a set of distinct promoters each specific to a certain vector design strategy: 1) the minP, allowing for packaging of larger transgenes but relying on enhancer strength and specificity, 2) the AFP promoter, a medium-sized cell-specific promoter to which enhancers add mainly strength, and 3) the PGK promoter, a ubiquitously active promoter to which enhancers can confer cell type specificity. It is conceivable that the large degree of promoter preference we observe is due to these distinct promoter types. To further explore the phenomenon of enhancer-promoter compatibility in the rAAV genome, future studies could determine the preference of enhancers for promoters of a similar type (e.g. different cell type-specific promoters). Nevertheless, we conclude that all types of promoters we have tested can be used with STARR-CRAAVT to identify promising enhancer candidates for regulatory element development.

Historically, it is a hallmark of enhancers that they can function independently of their relative position to the promoter inside a certain distance (Banerji et al. 1981; Gillies et al. 1983; Mercola et al. 1983; Thomas and Buecker 2023; Symmons et al. 2014; Zuin et al. 2022; Yang and Hansen 2024). In an episomal context, position independence has been confirmed by several STARR-Seq studies that validated identified enhancers by placing them upstream of a promoter in single luciferase assays (Arnold et al. 2013; Liu et al. 2017; Barakat et al. 2018). However, other studies using CapSTARR-Seq (Vanhille et al. 2015; Hussain et al. 2023) or exploring enhancer-promoter compatibility (Zabidi et al. 2015) validated identified enhancers only downstream of the promoter, similar to the positioning in the STARR-Seq setup. Indeed, when testing our STARR-CRAAVT-derived candidates, we find substantial differences in enhancer activity measured in luciferase assays dependent on the position in the rAAV expression cassette. Activity of three candidates was lost or reduced while activity of a fourth candidate increased ∼16-fold when moved from a downstream position upstream of the promoter. It appears possible that this position-sensitive enhancer activity is related to aspects of AAV genome biology, for example the oligomerization of episomal AAV genomes into concatemers (Yang et al. 1999; Dhungel et al. 2021). Indeed, a study comparing different plasmid- and lentivirus-based MPRA approaches found that enhancer activity can be influenced by relative positioning of the enhancer but also by other screening backbone design differences (Klein et al. 2020). Therefore, a broader systematic assessment of enhancer activity across different promoters and construct designs in the rAAV context is needed to better understand the observed position sensitivity.

In conclusion, our study demonstrates the successful integration of rAAV-based STARR-Seq and capture-based library generation into STARR-CRAAVT and infers guidelines for employing STARR-CRAAVT to identify enhancers for next-generation rAAV gene therapy vectors. Taking the observed promoter preference and position sensitivity into account, it is recommended to align the STARR-CRAAVT design as close as possible with the therapeutic rAAV vector design.

## METHODS

### *In-silico* library design

The library is based on the hepatocellular carcinoma (HCC) ATAC-Seq peaks downloaded from The Cancer Genome Atlas (TCGA; “LIHC_peakCalls.txt” file from the zip archive at https://api.gdc.cancer.gov/data/71ccfc55-b428-4a04-bb5a-227f7f3bf91c) and HCC H3K27ac active enhancer histone mark peaks (Hlady et al. 2019), downloaded from the NCBI Gene Expression Omnibus (GEO) database (Accession Number GSE112221) and converted to hg38 coordinates using the ‘liftOver’ tool from the UCSC genome bowser (Lee et al. 2022). HCC ATAC-Seq peaks overlapping transcription start sites taken from the GENCODE v37 human transcriptome (Frankish et al. 2021) were filtered out. The resulting peaks were intersected with the HCC H3K27ac peaks and only overlapping ATAC-Seq peaks were retained. To remove normal liver enhancers, human liver H3K27ac peaks from the ENCODE project (ENCODE Project Consortium et al. 2020) were downloaded and merged (Accession IDs: ENCFF275ZWJ, ENCFF218OOF, ENCFF193GDV, ENCFF042XXW, ENCFF976KNB, ENCFF805YRQ, ENCFF813EXR, ENCFF270BHR, ENCFF287VIA). Subsequently, HCC ATAC-Seq peaks overlapping these were excluded. In addition, the HCC ATAC-Seq peaks were supplemented with non-overlapping HCC H3K27ac peaks [Hlady et al., 2019] present in both replicates and filtered for normal human liver H3K27ac peaks as described above. Peaks larger than 1kb were split into 500 bp subregions separated by 300bp intervals, to cover the entire enhancer region. Finally, insulators were filtered out by removing peaks overlapping with constitutive CTCF peaks (downloaded from (Li et al. 2013)), and those overlapping microRNA genes (downloaded from miRbase v22 (Kozomara et al. 2019)) were also excluded. All peak intersections and filtering were performed using the ‘bedtools intersect’ tool from the BEDTools suite v2.26.0 (Quinlan and Hall 2010). The hg38 coordinates of the final candidate list were submitted to Agilent for generation of a custom SureSelect^XT^ target enrichment kit (Agilent Technologies, Santa Clara, CA, USA).

### Capture of the fragment library

Human DNA was isolated from blood samples of 9 healthy donors using the Genomic DNA kit-500/G (QIAGEN, Hilden, Germany). DNA was sheared using an ME220 ultrasonicator (Covaris, Woburn, USA) with an 8-microTUBE-130 AFA Fiber Strip V2 for 550 bp fragments. Sixteen micrograms of the fragmented products were size selected using SPRI beads (Beckman Coulter, Indianapolis, USA). The left-side selection protocol was utilized with a 0.65x ratio. After elution of size-selected fragments, 3 µg of DNA were subjected to the Agilent SureSelect^XT^ Target enrichment protocol including end-repairing, dA-tailing, ligation of the paired-end adaptor as well as amplification of the adaptor-ligated library. The quality of the amplified adaptor-library was assessed with a fragment analyser (Agilent Technologies, Santa Clara, CA, USA). The adaptor-ligated library was diluted to a concentration of 221 ng/µl and multiple hybridization reactions were set up. The hybridization of the DNA samples to the biotinylated probes was carried out according to the SureSelect^XT^ Target enrichment protocol for probe size > 3Mb. The hybridized samples were captured and purified by streptavidin-coated magnetic beads.

### STARR-CRAAVT screening plasmids

The minP (4.26) AAV-STARR-Seq plasmid has been described previously (Chan et al. 2023) and is based on the original STARR-Seq plasmid from Arnold et al. (Addgene #71509, (Arnold et al. 2013)). To create AFP and PGK AAV-STARR-Seq plasmids, AFP and PGK fragments were synthesized (GeneArt, Regensburg, Germany) and cloned into the minP AAV-STARR-Seq backbone digested with BglII and PspOMI.

### Library cloning

The captured fragment library was PCR amplified for in-fusion cloning into the vector backbone (forward primer: TAGAGCATGCACCGGACACTCTTTCCCTACACGACGCTCTTCCGATCT; reverse primer: GGCCGAATTCGTCGAGTGACTGGAGTTCAGACGTGTGCTCTTCCGATCT). The amplicons were purified using AMPure XP beads (Beckman Coulter, Indianapolis, USA). The STARR-CRAAVT backbones were restriction digested with AgeI and Sall. After purification, the amplified fragment library was ligated into the different STARR-CRAAVT backbones using the in-fusion cloning kit (Takara Bio, Shiga, Japan). The ligated DNA was precipitated and then transformed via electroporation to electrocompetent DH5a cells (Thermo Fisher Scientific, Waltham, USA).

### AAV production

AAV vectors were produced in CELLdiscs (Greiner Bio-One GmbH, Frickenhausen, Germany) by triple transfection of pRep/Cap6.2, pHelper, and either minP, AFP or PGK plasmids into HEK293-H (Thermo Fisher Scientific, Waltham, USA) cells using calcium phosphate. Purification was performed by polyethylene glycol precipitation, followed by iodixanol density gradient ultracentrifugation and ultrafiltration, as described previously (Strobel et al. 2019; Kuklik et al. 2021). Viral genomes were quantified by digital PCR using ITR-specific primers. Briefly, DNA was extracted from 25 µl of each AAV stock using ViralXpress Nucleic Acid Extraction Kit (Merck Millipore, Burlington, MA). Extracted DNA was diluted in a 10-step dilution and the QIAcuity Probe PCR Master Mix (QIAGEN, Hilden, Germany) and a 1× primer-probe mix for the detection of the AAV2 ITR (forward primer: GGAACCCCTAGTGATGGAGTT, reverse primer: CGGCCTCAGTGAGCGA, probe FAM-CACTCCCTCTCTGCGCGCTCG-MGB) (Sigma-Aldrich, Burlington, MA) was used according to the manufacturer’s instruction. dPCR reaction was performed on a QIAcuity One dPCR device (QIAGEN, Hilden, Germany).

### Cell culture

All cells used in this study were cultured at 37°C in a humidified atmosphere containing 5% CO2. Growth medium for HEK293-H, HepG2, and HaCaT cells consisted of Dulbecco’s modified Eagle’s medium (DMEM; Invitrogen Life Technology, Carlsbad, CA, USA) supplemented with 10% fetal bovine serum (FBS; Thermo Fisher Scientific, Waltham, USA). Cells were passaged using trypsin and maintained at sub-confluent densities.

### STARR-CRAAVT screen and next-generation sequencing

The STARR-CRAAVT screen was performed in three technical replicates, meaning that three wells per cell type were transduced and processed separately. HepG2 and HaCaT cells were grown to 80% confluency in 12 well plates. Cells were transduced with the STARR-CRAAVT AAV libraries using a multiplicity of infection (MOI) of 100,000 viral genomes per cell. The minP plasmid library was also transfected using 1 ug of DNA and Lipofectamine 3000 (Thermo Fisher Scientific, Waltham, USA). After incubation for 72h, cells were harvested using trypsin and DNA and RNA were extracted with an ALLprep 96 DNA/RNA kit (QIAGEN, Hilden, Germany). Prior to index PCR (iPCR), RNA samples were further processed as follows. Two micrograms of the purified total RNA were used for mRNA enrichment utilizing the NEBNext® Poly(A) mRNA Magnetic Isolation Module (New England Biolabs, Ipswich, USA), following manufacturer’s guidelines. The mRNA was eluted in a final volume of 13.5 µl Tris buffer and 11 µl of mRNA were used for cDNA synthesis using a gene-specific primer and SuperScript III reverse transcriptase (Thermo Fisher Scientific, Waltham, USA). The cDNA was then subjected to RNase A treatment (Thermo Fisher Scientific, Waltham, USA) for 1 h at 37 °C and purified using AMPure XP beads following manufacturer’s protocol. The candidate fragment cassette was amplified from the purified cDNA using exon-spanning primers and products were purified using AMPure XP beads. The concentration of the PCR amplicons was measured using a fragment analyser and 20 ng of PCR product as well as 20 ng of purified DNA were used for iPCR with UDI primers (New England Biolabs, Ipswich, USA). Additionally, 5 ng and 10 ng of plasmid and viral input DNA were used for iPCR, respectively. The iPCR products were purified twice using AMPure XP beads and pooled in an equimolar ration. The pooled samples were sequenced using a paired end read protocol (76+8+8+76) on a NextSeq (Illumina, San Diego, USA).

### STARR-CRAAVT analysis and candidate nomination

The NGS paired-end reads were mapped to the human genome (assembly hg38) using bowtie2 (v2.2.9) (Langmead and Salzberg 2012). Mapped reads were filtered for those mapping to a single genomic location using samtools (v1.2) (Li et al. 2009), and those mapping to the capture library target regions were quantified using the ‘featureCounts’ tool (Liao et al. 2014) of the Subread NGS alignment suite (v1.6.2) (Liao et al. 2013). Enhancer activity was determined by performing a differential expression analysis on the cell-extracted cDNA (mRNA) versus the cell-extracted DNA using the DESeq2 R package (v1.26) (Love et al. 2014, 2) with R (v3.10) (R Core Team 2021), with RNA/DNA ratio being used as a measure of enhancer strength. For fragment-level analyses, the fragments were processed using the BEDTools suite v2.26.0 (Quinlan and Hall 2010). First, they were extracted from the name-sorted paired-end BAM files (sorted with ‘samtools sort -n’) using the ‘bedtools bamtobed’ command with the ‘-bedpe’ flag, and their overall start and end coordinates were stored in the normal BED file format, with the mapping quality (MAPQ) placed in the score field and the read name in the name field. They were then filtered for MAPQ >= 30 using the Linux ‘awk’ command, their coordinates were rounded to the nearest 10 bp to account for reads with slightly trimmed ends, they were quantified using ‘bedtools groupby –c 4 –o count’, and finally they were intersected with the capture library target regions using ‘bedtools intersect’. For comparative STARR-CRAAVT analysis, processed sequencing results from the three technical replicates were filtered by removing candidates only detected in one replicate and then pooled. Correlations of STARR-CRAAVT activities (log2FC RNA/DNA) and analysis of enhancer overlaps (defined as log2FC RNA/DNA > 1 and adjusted p-value < 0.05) was done using R (v3.10). Candidates for validation were randomly chosen from a pool of enhancer candidates with the following criteria: Coefficient of variation below 0.5 and a minimum TPM count of 10.

### Luciferase assays

An AAV reporter backbone was generated (GeneArt, Regensburg, Germany) containing a Nano-Luciferase ORF driven by minP, AFP, or PGK flanked by AAV2 ITRs. To reduce transcriptional background, a set of polyA sites was included upstream of the promoters as described in Muerdter et al. (Muerdter et al. 2018). Fragments of enhancer candidates were PCR amplified from the libraries and cloned into the restriction-digested reporter plasmid using NEBuilder Hifi Assembly. Digestions were performed with EcoRV and SalI for upstream and downstream enhancer positioning, respectively.

HepG2 and HaCaT cells were grown to 80% confluency in a 96 well plate. One day after plating, 80 ng of plasmid was transfected using lipofectamine-3000 following the manufacturer’s protocol. Luciferase activity was detected 72h after transfection using the Nano-Glo® Luciferase Assay System according to the manufacturer’s instructions and measured with a SpectraMax i3x MiniMax 300 Imaging Cytometer (Molecular Devices, Sunnyvale, CA).

## ACKNOWLEDGMENTS

We thank Isabel Lang, Dragica Blazevic, Philipp Doll, Gudrun Zimmermann, Jenny Danner-Liskus, Christine Mayer, and Kai Zuckschwerdt for their technical assistance. Furthermore, we are grateful to the Boehringer AAV gene therapy team for the insightful discussions and feedback on the project. Also, we would like to thank Werner Rust, Alec Dick, and his team for high-throughput sequencing support. This research was funded by the Department of Eye Health & Research Beyond Borders at Boehringer Ingelheim Pharma GmbH & Co. KG, Biberach (Riss), Germany.

## AUTHOR CONTRIBUTIONS

According to CRediT: Conceptualization: R.B., P.C., M.O., S. Ketterer, J.P., S. Kreuz, U.M., C.S.; Formal analysis: R.B., M.O.; Funding acquisition: S. Kreuz, U.M., S.M.; Investigation: R.B., P.C., M.O., B.S.; Methodology: P.C., M.O., S.R., S. Kreuz, C.S.; Project administration: R.B., C.S.; Resources: B.S., J.P., S.M.; Supervision: U.M., S.M., C.S.; Visualization: R.B., Y.F.; Writing – original draft: R.B., C.S.; Writing – review & editing: R.B., M.O., Y.F., B.S., S. Ketterer, S.R., C.S.

