## Supplemental Figures for "STARR-CRAAVT: A platform to design cell type-specific regulatory elements for next-generation gene therapy"

### Supplemental figure S1

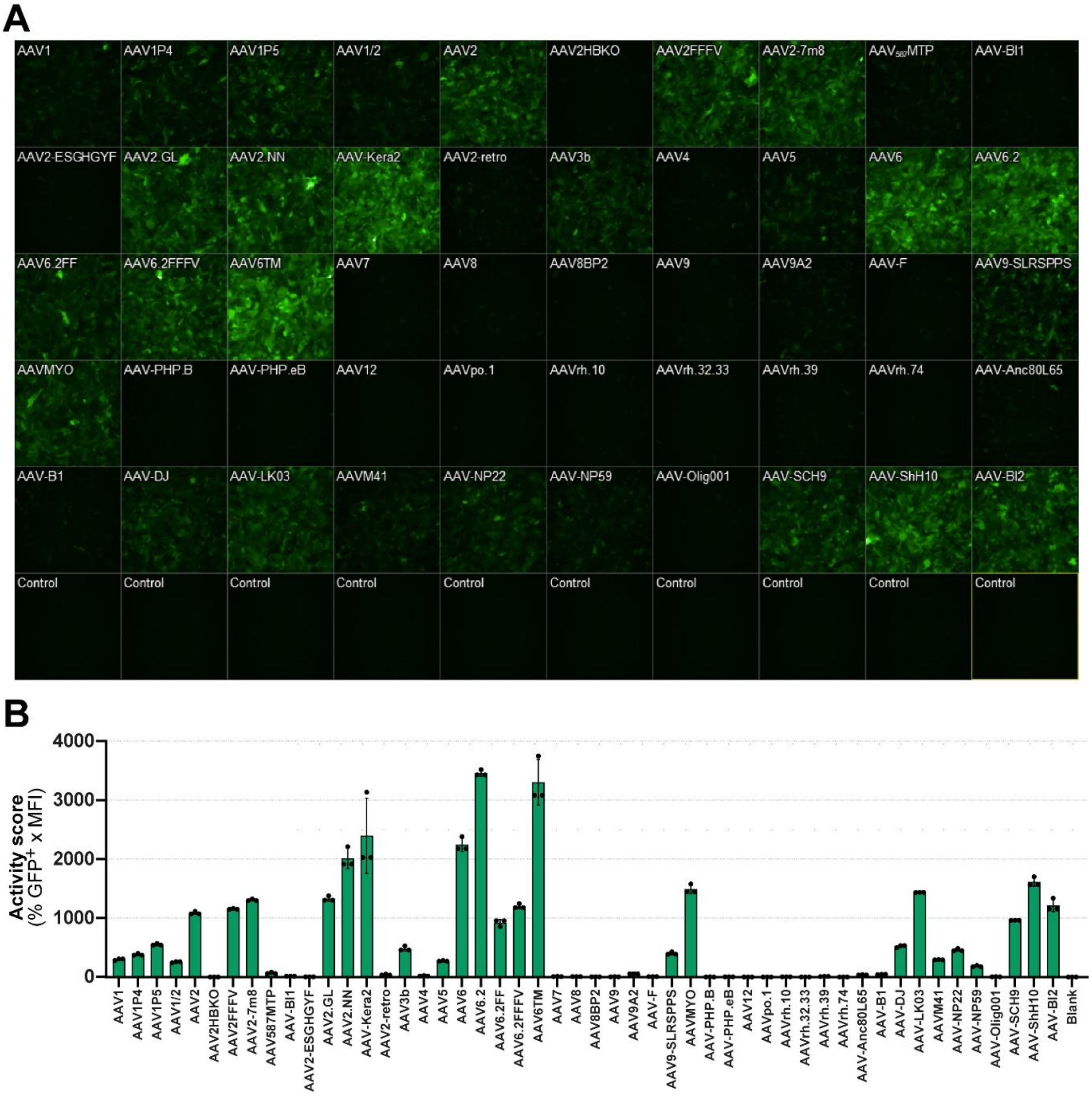


**Figure S1**. AAV6.2 transduces HaCaT cells. **(A)** HaCaT cells were subjected to a panel of AAV serotypes packaging an eGFP transgene (Weinmann *et al.*, 2022). **(B)** Quantification of (A) with the activity score being the product of the fraction of GFP-positive cells (% GFP+) and the mean fluorescent intensity (MFI).

### Supplemental figure S2

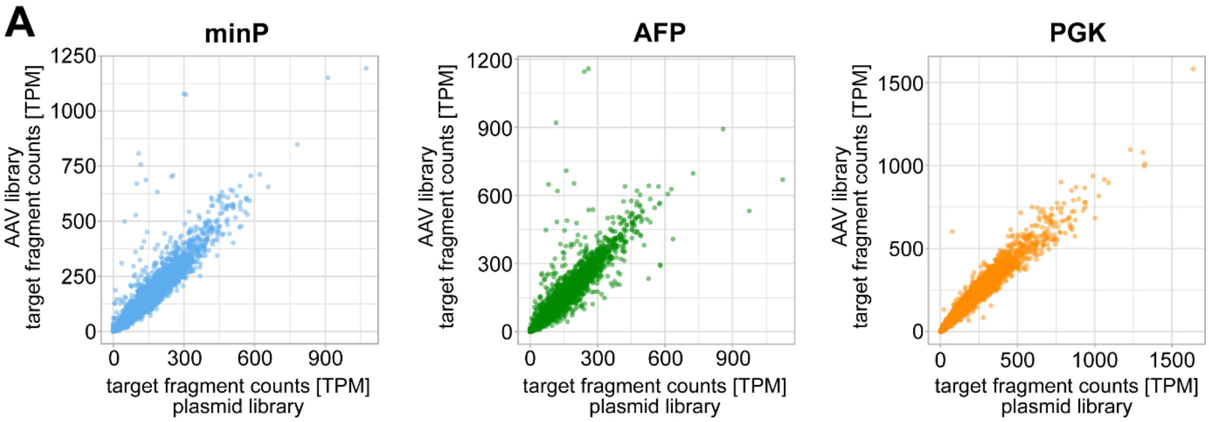


**Figure S2**. Analysis of STARR-CRAAVT candidate libraries after cloning and packaging. **(A)** Scatterplots comparing the fragment counts per candidate in the plasmid and the AAV libraries.

### Supplemental figure S3

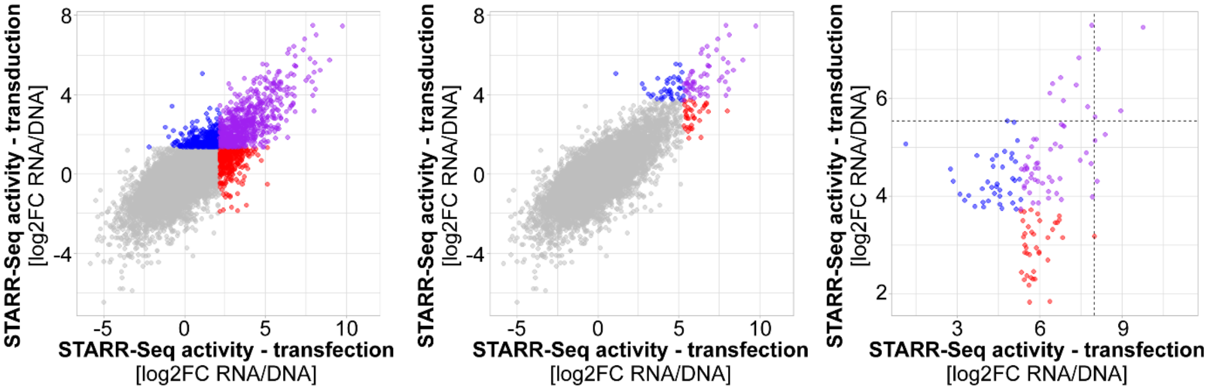


**Figure S3**. Effect of delivery mode on STARR-CRAAVT outcome. **Left and middle**: Scatterplots comparing STARR-Seq activity (measured as log2FC RNA/DNA) of minP candidate libraries between transfection and transduction of HepG2 cells. Graphs show only candidates for which a log2FC RNA/DNA could be calculated for both modalities. Colors indicate the top 1000 (left) or top 100 (middle) enhancer candidates that are common (●) or exclusive to transfected samples (●) or transduced samples (●). **Right:** Scatterplot comparing the STARR-Seq activity of the top 100 enhancers in HepG2 cells either transfected or transduced with the minP library. Colors indicate whether a candidate was found in the top 100 enhancers in transfected samples (●), transduced samples (●) or both (●). Dashed lines indicate the highest STARR-Seq activities of candidates exclusive to transfection and transduction.

### Supplemental figure S4

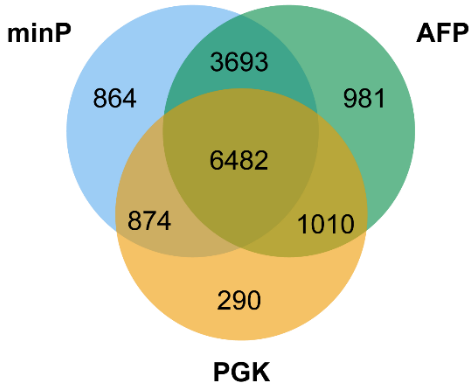


**Figure S4.** Venn diagram showing numbers of candidates covered by each input AAV library.
